# Using lifestyle information in polygenic modeling of blood pressure traits: a simple method to reduce bias

**DOI:** 10.1101/2024.06.05.597631

**Authors:** Francesco Tiezzi, Khushi Goda, Fabio Morgante

## Abstract

Complex traits are determined by the effects of multiple genetic variants, multiple environmental factors, and potentially their interaction. Predicting complex trait phenotypes from genotypes is a fundamental task in quantitative genetics that was pioneered in agricultural breeding for selection purposes. However, it has recently become important in human genetics. While prediction accuracy for some human complex traits is appreciable, this remains low for most traits. A promising way to improve prediction accuracy is by including not only genetic information but also environmental information in prediction models. However, environmental factors can, in turn, be genetically determined. This phenomenon gives rise to a correlation between the genetic and environmental components of the phenotype, which violates the assumption of independence between the genetic and environmental components of most statistical methods for polygenic modeling. In this work, we investigated the impact of including 27 lifestyle variables as well as genotype information (and their interaction) for predicting diastolic blood pressure, systolic blood pressure, and pulse pressure in older individuals in UK Biobank. The 27 lifestyle variables were included as either raw variables or adjusted by genetic and other non-genetic factors. The results show that including both lifestyle and genetic data improved prediction accuracy compared to using either piece of information alone. Both prediction accuracy and bias can improve substantially for some traits when the models account for the lifestyle variables after their proper adjustment. Our work confirms the utility of including environmental information in polygenic models of complex traits and highlights the importance of proper handling of the environmental variables.

**Author summary:** Many traits of medical relevance are “complex” in that they are affected by both genetic and environmental factors. Thus, using genetic and environmental information in statistical methods has the potential to increase the accuracy of phenotypic prediction, the ultimate goal of precision medicine. However, the correlation between the genetic and environmental components (that arises when environmental variables are themselves genetically determined) and the correlations between environmental measures can be problematic for most statistical methods used for modeling complex traits. In this work, we investigated these issues using 27 lifestyle measures in addition to genetic information for predicting diastolic blood pressure, systolic blood pressure, and pulse pressure in older individuals. We show that including lifestyle and genetic data resulted in more accurate predictions than either data type alone. Moreover, adjusting the lifestyle measures for the genetic and other non-genetic effects can help improve the predictions further.

## Introduction

Quantitative genetics theory posits that complex trait phenotypes (*P*) are affected by a genetic component (*G*) and an environmental component (*E*). These components result from the effect of many genetic variants and many environmental (or more precisely non-genetic) factors, respectively [1]. Thus, in its basic form, the quantitative genetic model can be written as

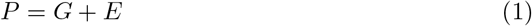

The genetic component can be further decomposed into additive (*A*), dominance (*D*), and epistatic (*I*) components, whereby *G* = *A* + *D* + *I* [1]. In this work, we are not concerned with dominance and epistasis, so we assume that *G* = *A* throughout, noting that this oversimplification does not invalidate our conclusions. Quantitative geneticists have been interested in partitioning the phenotypic variance 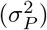 into genetic variance 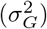 and environmental variance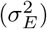, with the ratio 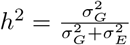 being the heritability of the trait [1]. (Because of the *G* = *A* assumption, narrow sense heritability and broad sense heritability are the same here).

In addition, the theory allows for the interaction between the genetic component and the environmental component through the genotype-by-environment interactions (*G × E*). From a genetic standpoint, this means that the effect of a genetic variant differs depending on the environmental factors [1]. Thus, the quantitative genetic model can be extended as

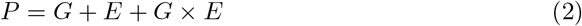

resulting in an additional variance component associated with genotype-by-environment interactions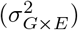.

Decomposing the phenotypic variance into its components and estimating the heritability of human complex traits has traditionally been done using pedigree information from close relatives such as twins [2]. However, with the advent of cheap genotyping, it has become possible to estimate the genomic or single nucleotide polymorphism (SNP) heritability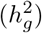, which is the proportion of phenotypic variance explained by regression on genotyped genetic variants using distantly related individuals [3]. Since the first application of the method to height [4], genomic heritability has been estimated for a plethora of complex traits and diseases, with estimates of 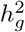 ranging from small to large [5, 6]. Among the many implications of genomic heritability is that it provides an upper bound to the accuracy of phenotypic prediction that a genomic-only model via polygenic scores (PGS) can achieve [7, 8]. (When prediction accuracy is measured as *R*^2^ – *i*.*e*., the squared correlation coefficient between true and predicted phenotypes –, then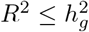). While recent advances in methods to compute PGS have provided much improved accuracies, these still have not reached estimates of 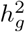 for the majority of traits [9–12].

One way to improve prediction accuracy is to include additional information in prediction models. Since complex traits are affected by genetic and non-genetic factors, incorporating the latter can potentially improve prediction accuracy. Lifestyle habits such as diet composition, physical activity levels, or smoking status are known to affect diseases and medically relevant traits such as diabetes, Alzheimer’s disease, and blood pressure [13–16]. Efforts to integrate lifestyle and genetic information for modeling complex trait variation have shown that both components explain a sizeable amount of the total phenotypic variance, with their relative importance being different depending on the trait [17, 18]. Zhou *et al*. (2021) also showed that adding lifestyle information improved the accuracy of phenotype prediction.

The inclusion of environmental information such as lifestyle opens to the possibility of modeling genotype-by-environment interactions. The importance of genotype-by-environment interactions for complex traits modeling and prediction has been documented thoroughly in agricultural species and model organisms [19–24]. Recent studies taking advantage of the large sample size of modern biobanks and consortia have also shown that genotype-by-environment interactions play a non-negligible role in the genetic architecture of human complex traits [25–30]. For example, Kerin and Marchini (2020) showed that about 9%, 4%, 2% and 12% of the variance of body mass index, systolic blood pressure, diastolic blood pressure, and pulse pressure, respectively, can be explained by genotype-by-environment interactions [27]. However, this has not generally translated into improved accuracy when including genotype-by-environment interactions into polygenic prediction models [18].

Environmental exposures can also be affected by genetic variants and become heritable. This phenomenon has been documented extensively in the behavioral sciences, as one way it may happen is indirectly through behavioral traits. For example, genetic variants affect whether a person is extroverted or introverted and the choice of social environment [31]. The result is that the genetic and environmental components in 2 become correlated, giving rise to genotype-environment correlation (*r*_*GE*_), which is not assumed to exist in the majority of polygenic models of complex traits. The presence of *r*_*GE*_ is problematic when modeling *G × E* because it induces “phantom” *G × G* (or epistasis) that can inflate estimates of *G × E*.

Thus, the presence of genotype-environment correlation and, additionally, correlations among environmental variables is problematic when modeling complex trait variation. While some attempts at accounting for these effects have been made [18], to the best of our knowledge, a thorough investigation of this topic is still missing. Thus, in this work, we investigated different ways of including genetic and environmental information (with and without their interaction) in statistical models to avoid bias from genotype-environment correlation on 1) variance components estimates and 2) the accuracy of phenotypic prediction. To do so, we used blood pressure traits (*i*.*e*., diastolic pressure, systolic pressure, and pulse pressure) and 27 lifestyle-related variables from UK Biobank [32] individuals, and methods and analytical strategies that have been successful in this type of study in agriculture.

## Materials and methods

### Data extraction and processing

The data used in this study were obtained from the UK Biobank (UKB) database. Data preparation and visualization of the results were performed in the R v.4.1.2 and 4.2.3 environment [33]. All plots were generated using the R package ggplot2 [34]. The phenotypes included were first-assessment readings of diastolic pressure (DP, UKB field: V4080-0.0) and systolic pressure (SP, UKB field: V4079-0.0). Data from manual measurements were not included in the present study. As in Tobin *et al*. [35] and Kerin and Marchini [27], measures of SP and DP were adjusted for individuals undergoing medication for blood pressure (UKB field: V6153-0.0, UKB field: V6177-0.0), adding 15 mmHg and 10 mmHg, respectively. Pulse pressure (PP) was calculated as the difference between the adjusted measures of SP and DP. Demographic variables included in the study were self-identified sex (SEX, UKB field: V31-0.0) and age at phenotyping (AGE, UKB field: V21003-0.0). The lifestyle component included 27 variables listed in Table 1. We note that some variables, such as body fat percentage, basal metabolic rate, and waist circumference, are not technically measures of lifestyle habits. However, since these measures are influenced by lifestyle factors (*e*.*g*., diet), we used them to capture some habits not explicitly modeled.

**Table 1.** Description of the 27 lifestyle variables.

| Description | Acronym | UK Biobank entry |
| --- | --- | --- |
| Townsend deprivation index at recruitment | Townsend | V189-0.0 |
| Time spent watching TV | TVtime | V1070-0.0 |
| Time spent using personal computer | PCtime | V1080-0.0 |
| Time spent driving | DRtime | V1090-0.0 |
| Sleep duration | sleep.d | V1160-0.0 |
| Ease of getting up in the morning | getup | V1170-0.0 |
| Smoking now | smoking.now | V1239-0.0 |
| Smoked in the past | smoked.past | V1249-0.0 |
| Cooked vegetables intake | veg.cook | V1289-0.0 |
| Oily fish intake | fish.oily | V1329-0.0 |
| Non-oily fish intake | fish.lean | V1339-0.0 |
| Processed meat intake | meat.proc | V1349-0.0 |
| Poultry intake | poultry | V1359-0.0 |
| Beef intake | beef | V1369-0.0 |
| Lamb intake | lamb | V1379-0.0 |
| Pork intake | pork | V1389-0.0 |
| Cheese intake | cheese | V1408-0.0 |
| Salt added to food | salt | V1478-0.0 |
| Tea consumption | tea | V1488-0.0 |
| Coffee consumption | coffee | V1498-0.0 |
| Alcohol intake frequency | alc1 | V1558-0.0 |
| Body fat percentage | BFP | V23099-0.0 |
| Basal metabolic rate | BMR | V23105-0.0 |
| Waist circumference | waist | V48-0.0 |
| Number of days per week walked 10 minutes | walk.d | V864-0.0 |
| Number of days per per week of vigorous physical | act0.d | V884-0.0 |
| Week of moderate physical activity | act1.d | V904-0.0 |

Only individuals belonging to ‘White British’ ethnicity (UKB field: V21000-0.0) and ‘Caucasian’ genetic ethnic grouping (UKB field: V22006-0.0) that had complete records (*i*.*e*., no missing values) for all the variables above were retained for the analyses. In this study, we used the array genotype data and processed them to retain genetic variants with Minor allele frequency (MAF) ≥ 0.01, Minor Allele Count (MAC) ≥ 5, genotype missing rate *<* 0.1 and Hardy-Weinberg Equilibrium (HWE) p-value *>* 1e-10. We also pruned the data for relatedness, such that no pairs of retained individuals had genomic relationships greater than 0.025. This filtering was done using PLINK 1.9. After the editing described, *n* = 98, 676 individuals, *p* = 595, 948 genetic variants, and *c* = 27 lifestyle variables were available for subsequent analysis. Using demographic variables SEX and AGE, eight cohort groups (COH) were created following the representation in Figure 4b of Kerin and Marchini [27]. This involved creating four age groups with break points at 51, 58 and 63 years, then concatenating the age group information to the SEX variable. This led to having classes denoted as 0-1 (female, age from 40 to 51, 12,934 individuals), 0-2 (female, age from 52 to 58, 11,497 individuals), 0-3 (female, age from 59 to 63, 10,962 individuals), 0-4 (female, age from 64 to 70, 12,975 individuals), 1-1 (male, age from 40 to 51, 13,352 individuals), 1-2 (male, age from 52 to 58, 11,395 individuals), 1-3 (male, age from 59 to 63, 11,384 individuals), and 1-4 (male, age from 64 to 70, 14,177 individuals). The eight groups showed similar representation in the final dataset. In addition, eight random groups (RND) with the same number of individuals as in COH were created by randomly assigning each individual to a group.

Before the statistical analysis, all the response variables (*i*.*e*., blood pressure traits and lifestyle variables when treated as phenotypes for variance decomposition) were centered to mean equal to 100 and standard deviation equal to 10, the continuous and ordinal lifestyle variables were centered to mean equal to 0 and standard deviation equal to 1.

### Statistical analysis

The statistical analysis involved fitting linear mixed models (LMM) to partition the phenotypic variance into residual, genetic, lifestyle, and genotype-by-lifestyle interaction components and make predictions for yet-to-be-observed phenotypes. The COH variable was added to the models to disentangle the impact of sex and age (at phenotyping) from the actual lifestyle component.

#### Partitioning variance of lifestyle variables

We started by partitioning the variance of each of the 27 lifestyle variables into demographic, genetic, lifestyle and residual components. Provided that **M** is a *n × c* matrix containing the 27 (raw) lifestyle variables (centered and scaled), the model used for this purpose was:

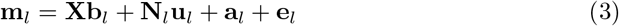

Where **m**_*l*_ is the *l*^*th*^ column of the **M** matrix (lifestyle variable of interest); **X** is an *n × q* incidence matrix for the intercept and COH; **b**_*l*_ is a *q*-vector of their fixed effects; **N**_*l*_ is a modification of the **M** matrix where the *l*^*th*^ column is replaced with a vector of random values (sampled from a Normal distribution with mean equal to 0 and standard deviation equal to 1); **u**_*l*_ is a *c*-vector of effects (regression coefficients) for the other lifestyle variables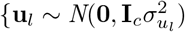, **I**_*c*_ is a *c × c* identity matrix}; **a**_*l*_ is a *n*-vector of individual additive genetic effects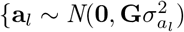, with **G** being a SNP-derived genomic relationship matrix (GRM) among the individuals [4] computed using GCTA [36]}; **e**_*l*_ is a n-vector of residuals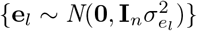.

Using the estimated effects from this model, two matrices of “adjusted” lifestyle variables were obtained. First, **E** is a *n × c* matrix where each column is defined as **e**_*l*_. Hence, **E** is a matrix containing residual values (adjusted for COH, genetics and other lifestyle variables) from (3) applied to each lifestyle variable. Second, **L** is a *n × c* matrix where each column is defined as **N**_*l*_**u**_*l*_. Hence, **L** is a matrix that contains lifestyle values for each variable predicted using the other lifestyle variables and adjusted for COH and genetics.

#### Modeling blood pressure traits

After partitioning the variance of the lifestyle variables and collecting information in the **M, L** and **E** matrices, we sought to model the three blood pressure traits.

A baseline model (ℳ_0_) accounting for the effects of cohort and additive genetic effects was defined as:

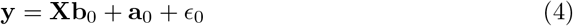

Three models (ℳ_1_, ℳ_2_, ℳ_3_) accounting for the effects of cohort and the three definitions of lifestyle variables, respectively, were defined as:

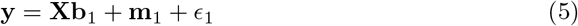

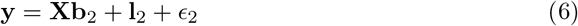

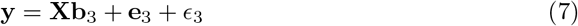

Three models (ℳ_11_, ℳ_12_, ℳ_13_) accounting for the effects of cohort, the three definitions of lifestyle variables, respectively, and additive genetic effects were defined as:

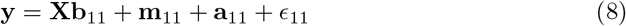

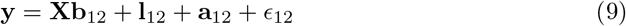

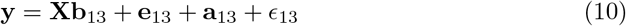

Three models (ℳ_21_, ℳ_22_, ℳ_23_) accounting for the effects of cohort, the three definitions of lifestyle variables, respectively, the additive genetic effects as well their interaction were defined as:

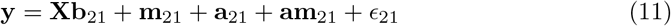

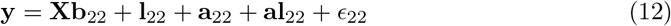

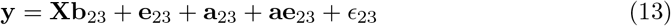

For each model ℳ_*t*_, the relevant terms are defined as follows. **X** is an *n × q* incidence matrix for the intercept and COH; **b**_*t*_ is a *q*-vector of their fixed effects; **a**_*t*_ is a *n*-vector of individual additive genetic effects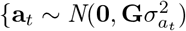, with **G** being a SNP-derived genomic relationship matrix}; **m**_*t*_ is a *n*-vector of individual raw lifestyle effects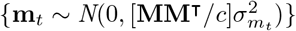; **l**_*t*_ is a *n*-vector of individual ‘predicted’ lifestyle effects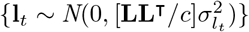; **e**_*t*_ is a *n*-vector of individual adjusted lifestyle effects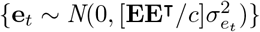; **am**_*t*_ is a *n*-vector of individual interaction effects between additive genetic effects and raw lifestyle effects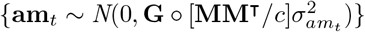; **al**_*t*_ is a *n*-vector of individual interaction effects between additive genetic effects and ‘predicted’ lifestyle effects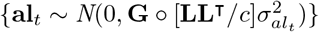; **ae**_*t*_ is a *n*-vector of individual interaction effects between additive genetic effects and adjusted lifestyle effects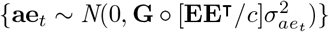; *ϵ*_*t*_ is a n-vector of residuals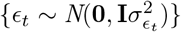. Note that º denotes the Hadamard product.

To check the impact of population structure, we implemented the following model (ℳ_01_):

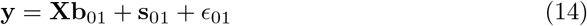

where **s**_01_ is a *n*-vector of their effects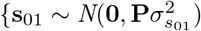, **P** is *n × n* ‘truncated’ GRM based on the first *r* = 20 principal components of **G**}

For the RND split, we also implemented a model (ℳ_02_) with only the cohort effect:

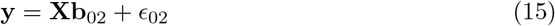

#### Model fitting

The models were implemented in a Bayesian framework using the R package BGLR v.1.1.0 [37]. The “fixed” effects were assigned bounded uniform priors and the variance components were assigned inverted *χ*^2^ priors [37]. The models were implemented running a total of 60,000 iterations, discarding the first 10,000 as burn-in, thinning every 50 iterations, and other default parameters.

#### Inference on variance components

To decompose the variance of the phenotypic traits (SP, DP and PP), all the models were implemented on the whole dataset (n=98,676). The variance explained by each of the effects was calculated at each iteration of the Gibbs sampler as follows: for the cohort effect, Var(**Xb**_*t*_); for the additive genetic effects, Var(**a**_*t*_); for the raw lifestyle effects, Var(**m**_*t*_); for the adjusted lifestyle effects, Var(**e**_*t*_); for the ‘predicted’ lifestyle effects, Var(**l**_*t*_); for the interaction between additive genetic effects and raw lifestyle effects, Var(**am**_*t*_); for the interaction between additive genetic effects and adjusted lifestyle effects, Var(**ae**_*t*_); for the interaction between additive genetic effects and ‘predicted’ lifestyle effects, Var(**al**_*t*_).

Similarly, to decompose the variance of the 27 lifestyle variables, the proportion of variance explained by each effect was calculated at each iteration of the Gibbs sampler as follows: for the cohort effect, Var(**Xb**_*l*_); for the lifestyle effects, Var(**N**_*l*_**u**_*l*_); for the additive genetic effects, Var(**a**_*l*_). The model for the decomposition of the lifestyle variables was still implemented on the whole dataset. This is due to the necessity to obtain values for the **E** and **L** matrices for both training and validation sets.

The mean of the posterior distribution for the variance explained was used as a point estimate and 95% empirical credible interval was used as a measure of error. These were calculated using the R package ‘TeachingDemos’ [38]. We note that because the response variables were centered to have mean equal to 100 and standard deviation equal to 10, these variances can be interpreted as percentages.

#### Estimation of genetic correlations between age groups

We estimated genetic correlations (*r*_*G*_) between age groups for each trait. We defined two groups – YOUNG (Y), which included cohorts 0-1, 0-2, 1-1, 1-2, and OLD (O), which included cohorts 0-3, 0-4, 1-3, 1-4. For each blood pressure phenotype, we fitted a bivariate GREML model implemented in GCTA v.1.94.1 [36], treating the phenotypes of individuals in the two groups as two different traits. We included the top ten principal components of the GRM as covariates. We computed approximate 95% confidence intervals as 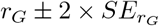 where 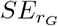 is the standard error of the genetic correlation estimate.

#### Prediction of blood pressure measures in older individuals

All the models were also implemented to predict blood pressure measures as manifested in individuals recruited at older ages (Y-O prediction). The individuals were split into training and validation sets, with individuals in the Y group used as the training set and individuals in the O group used as the validation set. The prediction models yielded predicted values (**ŷ**) for the individuals in the validation set, which were used to obtain measures of the models’ predictive ability. There were:

*Accuracy*, defined as the Pearson correlation coefficient between observed and predicted values:

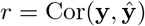

*Bias*, defined as the coefficients of a simple linear regression of the observed values on the predicted values, i.e.:

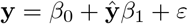

The intercept parameter *β*_0_ was used as a measure of systematic overestimation or underestimation of the predicted values, its expectation for unbiased predictions being equal to 0. The slope parameter *β*_1_ was used as a measure of overestimation or underestimation of the predicted values depending on the distance from the mean, with its expectation for unbiased estimates being equal to 1 [39]. The results obtained with the Y-O prediction were compared to a cross-validation in which the dataset was randomly split into training and prediction sets, with all cohorts equally represented. This was obtained using the RND groups described above.

## Results

### Variance components for lifestyle variables

Results for the partition of variance for the lifestyle variables are reported in Fig 1 and S1 Table..

**Fig 1.**
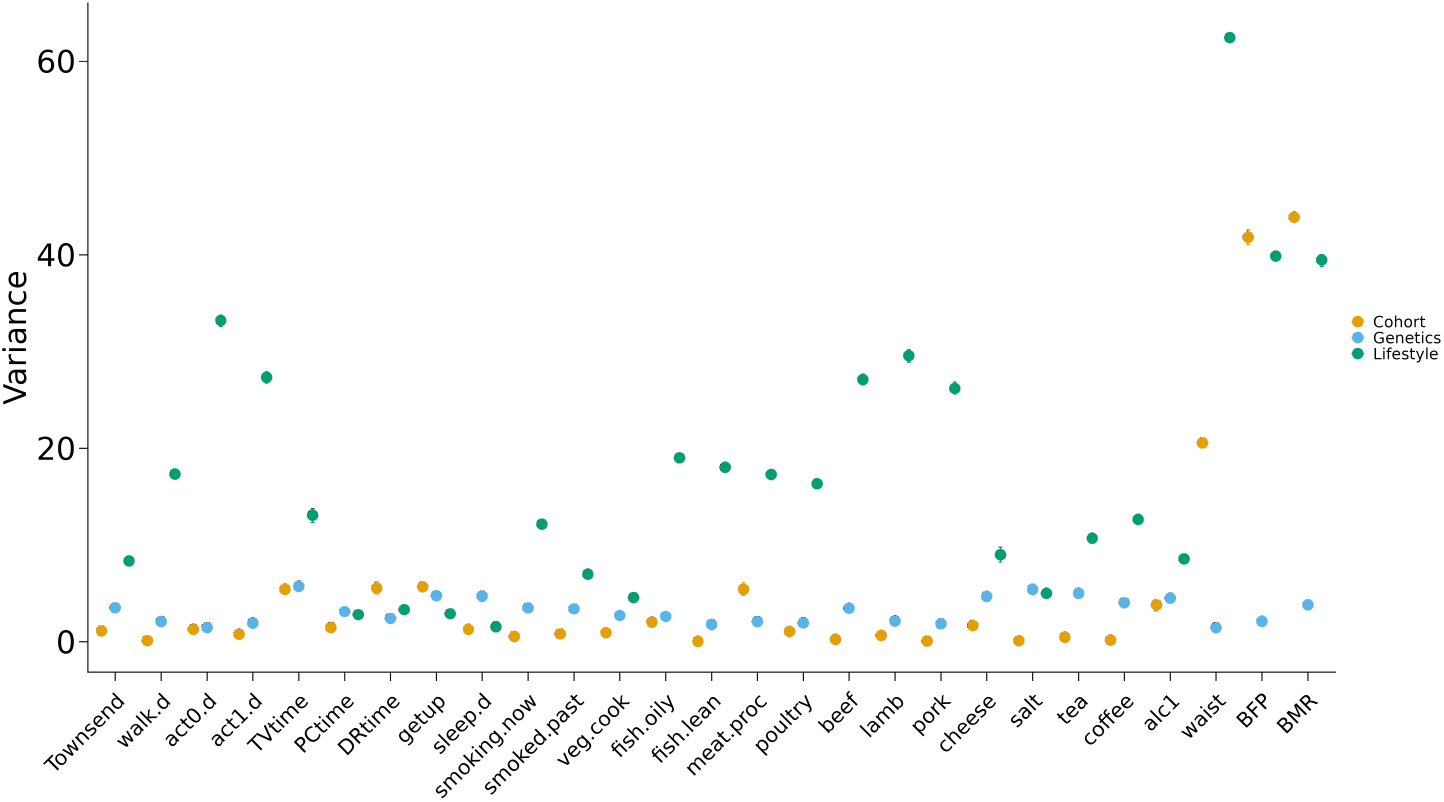
Variance components estimates for the 27 lifestyle variables. Bars represent 95% empirical credible interval for the posterior distribution of the parameters.

For each lifestyle variable, the effect with the strongest impact was the one of the other 26 lifestyle variables, which explained 18% of total variance on average across lifestyle variables, ranging from 1.5% for sleep duration to 62.5% for waist circumference. The second in order of magnitude was the effect of the cohort, which explained 5% of total variance on average across lifestyle variables. The variables for which the effect of cohort was largest (explaining more than 10% of the total variance) were those related to body size and physiology (*i*.*e*., basal metabolic rate, body fat percentage, and waist circumference). On the other hand, the variables for which the effect of cohort was smallest (explaining less than 1% of the total variance) were those related to eating habits and smoking.

The proportion of variance explained by the additive genetic effects (*i*.*e*., genomic heritability 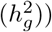 was 3% on average over all variables. Physical activity-related variables as well as lean fish, pork, and poultry consumption were the variables with the smallest 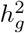 (less than 2%), while TV time, tea consumption, salt consumption showed 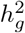 larger than 5%.

The residual component was the largest for all the lifestyle variables (ranging from 65% to 93%), except for basal metabolic rate, body fat percentage and waist circumference, for which it explained less than 20% of the total variance.

### Dimensionality of the lifestyle kernels

Results from the eigenvalue decomposition of the three lifestyle kernels used are reported in Table 2.

**Table 2.**
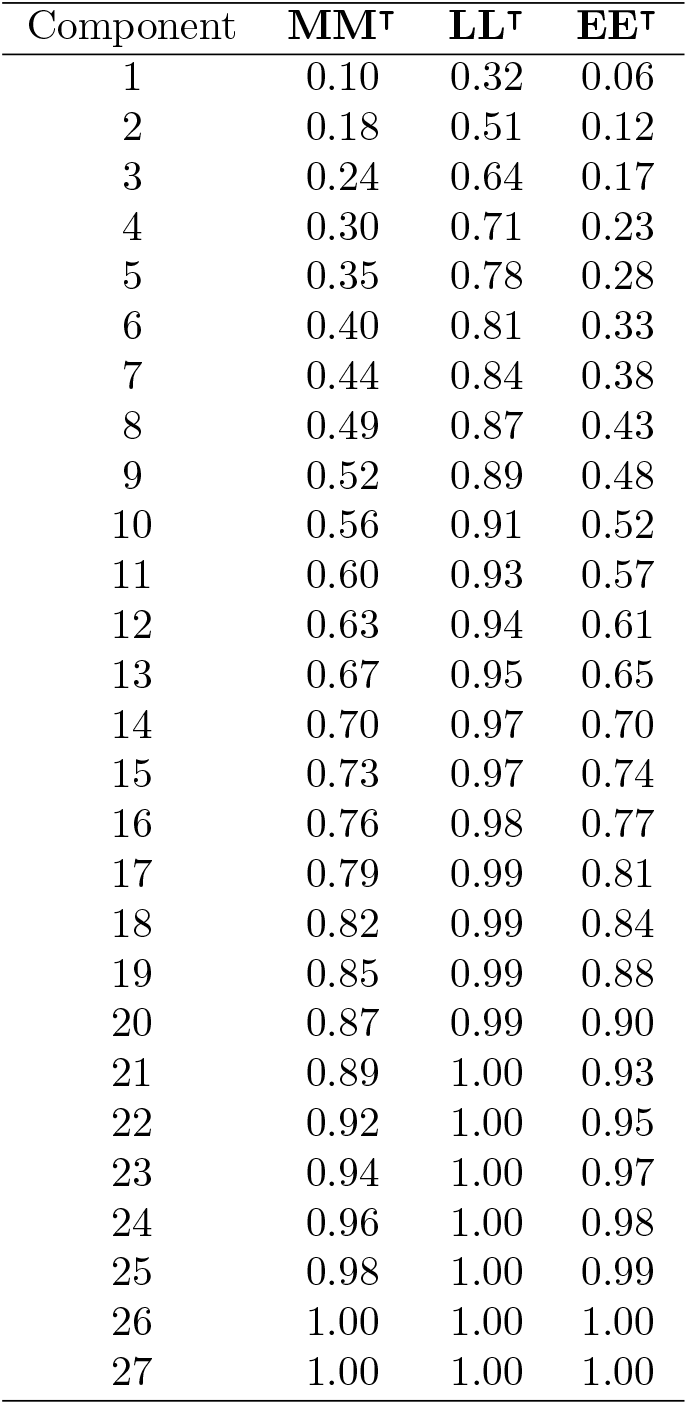
Proportion of variance explained (cumulative) by the components of the three lifestyle kernels, following eigenvalue decomposition. **MM**^⊺^ represents a kernel built on the 27 raw lifestyle variables. **LL**^⊺^ represents a kernel built on the 27 lifestyle variables adjusted by cohort, additive genetic and residual effects. **EE**^⊺^ represents a kernel built using the 27 lifestyle variables adjusted for the effects of cohort, additive genetics and the other lifestyle variables.

The **MM**^⊺^ kernel was built using the raw lifestyle variables. The decomposition of this kernel shows some degree of collinearity among the 27 lifestyle variables, with the first component explaining almost 10% of the variance and the first 9 components explaining 50% of the variance. On the other hand, the **LL**^⊺^ kernel was built on marginal predictions (after adjusting for the effects of cohort and additive genetics) of each lifestyle variable using the other lifestyle variables as predictors. Consequently, this kernel showed reduced dimensionality, with the first component explaining more than 30% of the variance and the first 9 components explaining 90% of the variance. The **EE**^⊺^ kernel was constructed using residual deviations after adjusting for the effects of cohort, additive genetics, and other lifestyle variables. This kernel showed similar dimensionality to the **MM**^⊺^ kernel, with the first component explaining 6% of its variance and the first 10 components explaining 50% of its variance.

### Variance components for blood pressure traits

Fig 2 and S2 Table. show the phenotypic variance partition for the three blood pressure traits. Each trait showed a different pattern of the contribution of the different effects to the total variation.

**Fig 2.**
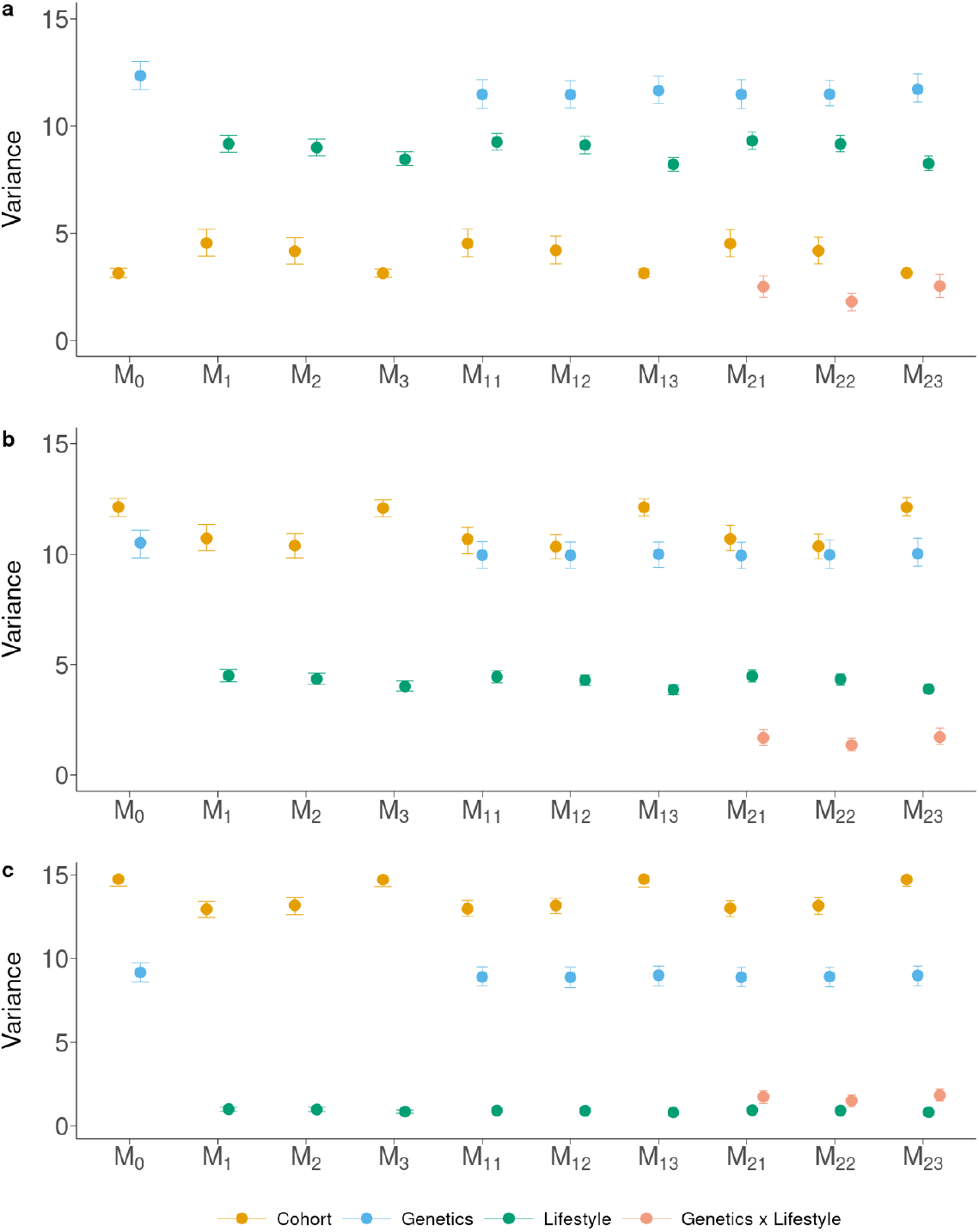
Variance components estimates for a) diastolic pressure, b) systolic pressure, and c) pulse pressure. Bars represent 95% empirical credible interval for the posterior distribution of the parameters.

The cohort component showed different trends depending on the trait, whereas it was consistent for each trait across the models. Having a stable partition despite changes in model specification suggests that the genetic and lifestyle components were well balanced over the cohorts, *i*.*e*., adding effects to the model did not change the estimates of the variance components for the existing effects. For DP, the cohort effect explained 3.13% of the variance in ℳ_0_, similarly to all the other models when the **EE**^⊺^ matrix was used as Lifestyle kernel (ℳ_3_, ℳ_13_, ℳ_23_), while the estimates were larger when the **MM**^⊺^ and **LL**^⊺^ kernels were used (around 4.5% for ℳ_1_, ℳ_11_, ℳ_21_, and 4.2% for ℳ_2_, ℳ_12_, ℳ_22_) regardless of the other effects in the model. For SP the pattern was inverted, with 12.1% of the variance explained by the cohort effect for ℳ_0_, ℳ_3_, ℳ_13_, and ℳ_23_, while the other models showed slightly smaller variance explained by this effect (between 10.4% and 10.7%). Similar pattern was found for PP, where this component was larger for ℳ_0_, ℳ_3_, ℳ_13_, and ℳ_23_ (~14.7% of the variance) and smaller for the other models (~13.1% of the variance). The genetic component variance showed stable estimates across all models for the three traits. Estimates were slightly larger with ℳ_0_ (12.3%, 10.5% and 9.2% for DP, SP and PP, respectively) than with the other models (11.5%, 10.0% and 9.0% for DP, SP and PP, respectively). However, the credible intervals for the point estimates overlapped across models. The lifestyle component, regardless of the kernel used, showed stable estimates across models, explaining less than 10% of the variance for DP, less than 5% for SP and about 1% for PP. The raw variables (*i*.*e*., **MM**^⊺^) tended to explain slightly more variance than the adjusted ones. Among the latter, the **LL**^⊺^ kernel explained more variance than the **EE**^⊺^ kernel. The genotype-by-lifestyle interaction component was consistent across traits and models, yet some differences could be found. The genotype-by-lifestyle interaction component explained 2.5%, 1.8% and 2.5% of the total phenotypic variance for ℳ_21_, ℳ_22_ and ℳ_23_, respectively, for DP, 1.8%, 1.5% and 1.8% for ℳ_21_, ℳ_22_ and ℳ_23_, respectively, for PP, and 1.7%, 1.4% and 1.7% for ℳ_21_, ℳ_22_ and ℳ_23_. The model using the **LL**^⊺^ kernel (ℳ_22_) provided smaller estimates of the genotype-by-lifestyle interaction component than the other kernels.

Estimates of residual variance were generally larger for models that did not include the genetic component, such as ℳ_1_, ℳ_2_, ℳ_3_. The residual variance in these models was around 88%, 84%, and 84% for DP, SP and PP, respectively. The pattern for the other models depended on the trait. In general, the residual variance tended to be larger for ℳ_0_ because it did not include the lifestyle component. However, the differences with the other models were small for PP and SP, and slightly larger for DP.

Also, for SP the residual variance was larger when the lifestyle variance decreased. The estimate of the genotype-by-lifestyle interaction component was inversely related to the estimate of the residual variance, showing that the interaction, when included, explained residual variance rather than variance from other effects.

### Predictive ability for blood pressure traits in older individuals

Predicting phenotypes across age groups is a difficult task since [27] showed that an environmental score built using similar lifestyle variables to the ones used in this study had a different distribution among the eight cohort groups. In addition, Fig 3 shows that the genetic correlation across age groups is significantly different from 1 for the three blood pressure traits, with lower estimated value for PP (*r*_*G*_ = 0.82) and higher values for DP (*r*_*G*_ = 0.91) and SP (*r*_*G*_ = 0.90). This suggests a different genetic architecture between younger and older individuals.

**Fig 3.**
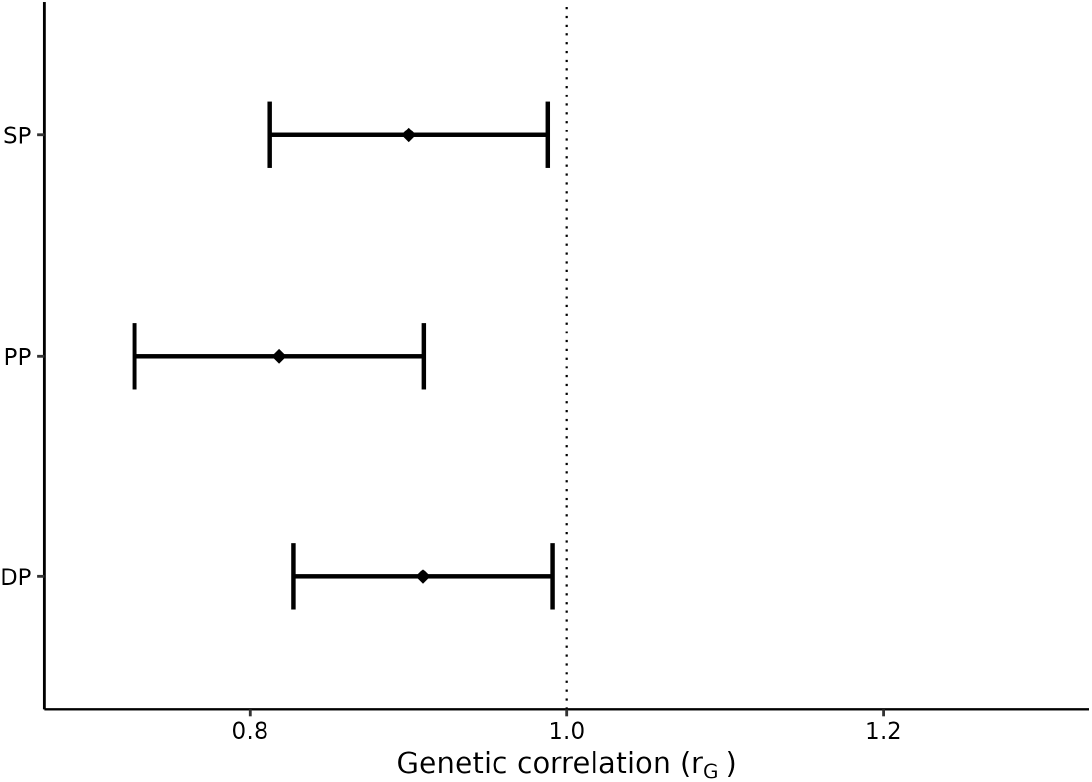
Genetic correlation between YOUNG (age 40-58) and OLD (age 59-70) groups for diastolic pressure, systolic pressure, and pulse pressure. Bars represent an approximate 95% confidence interval for the point estimate.

Fig 4 and S3 Table. show the accuracy of prediction obtained with the different models when training the models using the younger individuals and validating on the older ones, with both sexes being included.

**Fig 4.**
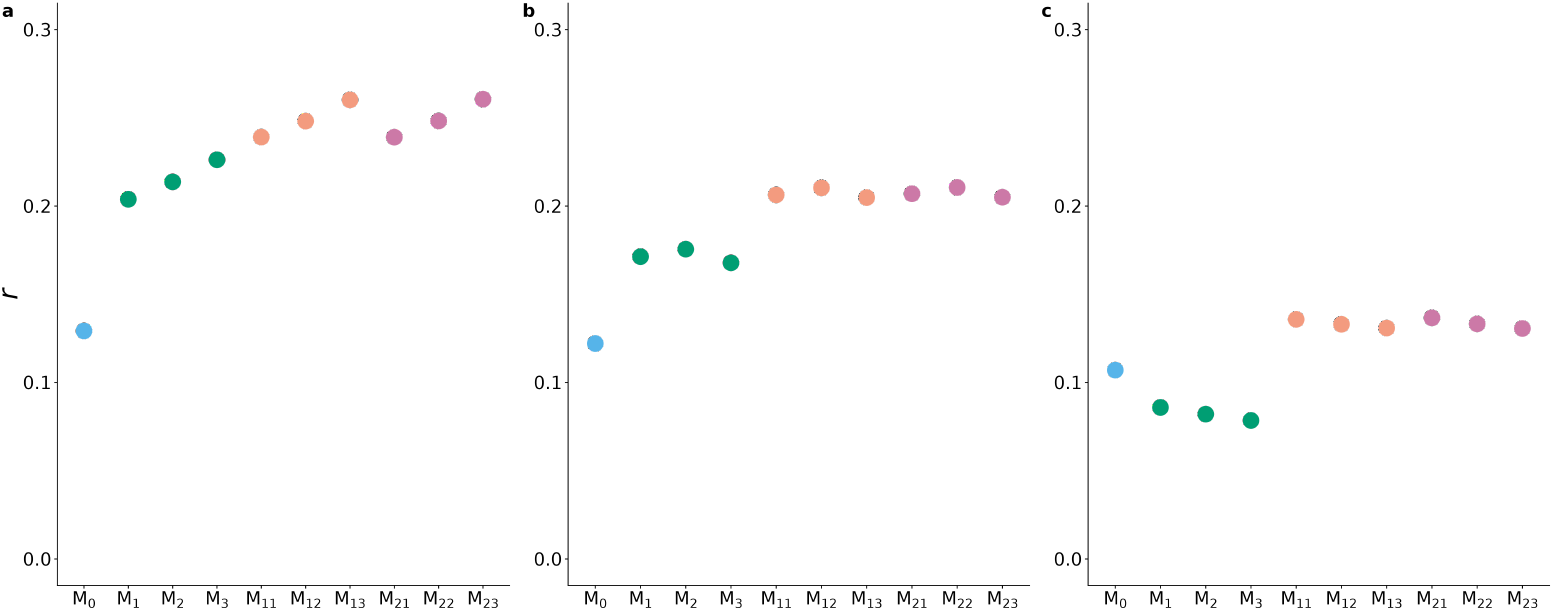
Accuracy of prediction measured as the correlation coefficient (*r*) between true phenotypes and predicted phenotype for a) diastolic pressure, b) systolic pressure and c) pulse pressure. The colors indicate the broad class of models, *i*.*e*., including genetic effects only (light blue), lifestyle effects only (green), genetic and lifestyle effects (orange), and genetic, lifestyle and interaction effects (purple).

In Fig 4, DP and SP show a similar pattern. Here, ℳ_0_ provided the lowest prediction accuracy (around 0.12) for both traits. The models that included only the lifestyle variables, whether raw or adjusted, (*ℳ*_1_, *ℳ*_2_, *ℳ*_3_) led to increased accuracy, which was between 0.20 and 0.23 for DP, and 0.16 and 0.18 for SP. The inclusion of both genetic and lifestyle effects (*ℳ*_11_, *ℳ*_12_, *ℳ*_13_) further increased the accuracy of prediction to 0.24-0.26 for DP and 0.20-0.21 for SP. The inclusion of the interaction (*ℳ*_21_, *ℳ*_22_, *ℳ*_23_) did not change the predictive ability. For PP, the pattern was different and saw a decrease in prediction accuracy when replacing the genetic effect with the lifestyle effects (from 0.11 for *ℳ*_0_ to 0.08 for *ℳ*_1_, *ℳ*_2_, *ℳ*_3_). The inclusion of both the effects (*ℳ*_11_, *ℳ*_12_, *ℳ*_13_) increased the prediction accuracy to around 0.13. The inclusion of the interaction (*ℳ*_21_, *ℳ*_22_, *ℳ*_23_) did not provide any increase in accuracy.

When considering the different sets (*i*.*e*., raw, adjusted, or ‘predicted’) of lifestyle variables, only in the case of DP there were substantial differences among the models. Accuracy was lowest when using the raw variables (*ℳ*_1_, *ℳ*_11_, and *ℳ*_21_), intermediate when using the adjusted variables (*ℳ*_2_, *ℳ*_12_, and *ℳ*_22_), and highest when using the ‘predicted’ variables (*ℳ*_3_, *ℳ*_13_, and *ℳ*_23_), with a difference in accuracy of about 0.01 among models including the different definitions of lifestyle variables. No appreciable differences were found when comparing the different sets of lifestyle variables for SP and PP.

As expected given our aggressive filtering to obtain a genetically homogeneous population as possible, the model including the first 20 principal components of the GRM (*ℳ*_01_) provided negligible prediction accuracy (*r* = 0.01 − 0.03) for all the traits (S3 Table.).

From the same cross-validation, Fig 5 and S3 Table. report the bias of the predictions as the intercept and slope, respectively, of the regression of the observed phenotypes on the predicted phenotypes.

**Fig 5.**
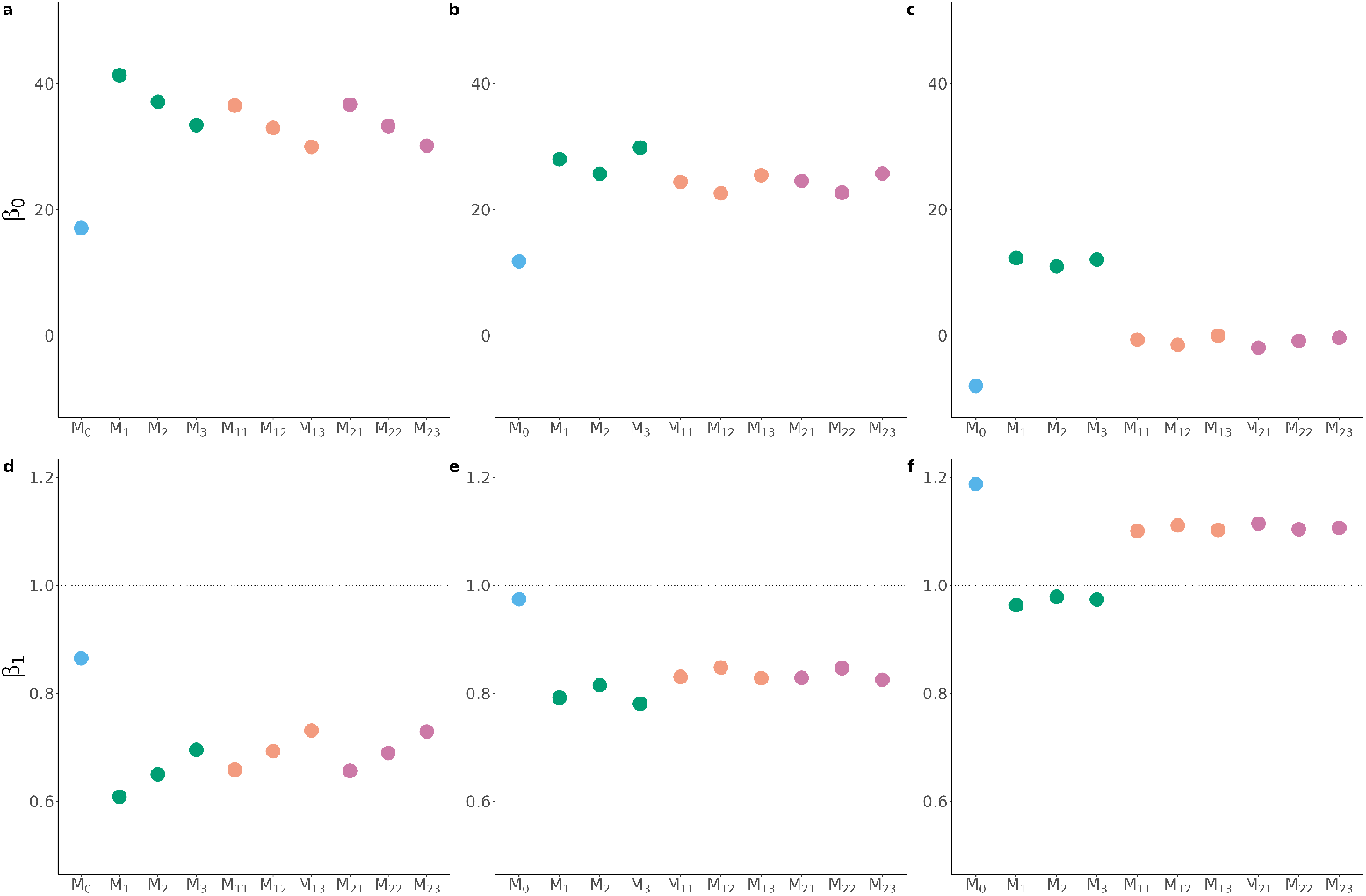
Bias of predictions measured as the intercept [*β*_0_; panels a), b) and c) for diastolic pressure, systolic pressure, and pulse pressure, respectively] and slope [*β*_1_; panels d), e) and f) for diastolic pressure, systolic pressure, and pulse pressure, respectively] of the regression of the true phenotypes on the predicted phenotypes. The colors indicate the broad class of models, *i*.*e*., including genetic effects only (light blue), lifestyle effects only (green), genetic and lifestyle effects (orange), and genetic, lifestyle and interaction effects (purple). The dotted lines indicate the expected values of *β*_0_ and *β*_1_ for unbiased predictions.

In Fig 5a,b,c, the suggestive horizontal line at 0 shows the optimal value for the estimate of the regression intercept. Here, all of the models provided an overestimation of DP and SP values, however, there were differences between the models. *ℳ*_0_ yielded the estimates closest to 0 for DP and SP (0.17 and 0.12 for DP and SP, respectively). *ℳ*_1_, *ℳ*_2_, *ℳ*_3_ consistently gave higher inflation, with values of 41, 37 and 33, respectively, for DP, and 28, 26 and 29, respectively, for SP. *ℳ*_11_, *ℳ*_12_, *ℳ*_13_ were able to reduce this inflation, reporting intercept values of 37, 33 and 30, respectively, for DP, and values of 24, 23 and 25, respectively, for SP. *ℳ*_21_, *ℳ*_22_, *ℳ*_23_, which included the interaction term, gave intercept estimates very similar to the models without such interaction. PP showed a completely different pattern. In particular, models including both additive genetic effects and lifestyle effects (with or without the interaction between these two components) provided unbiased predictions (*i*.*e*., intercept values very close to 0). On the other hand, the model including only additive genetic effects (*ℳ*_0_) underestimated PP values, while models including only lifestyle variables (*ℳ*_1_, *ℳ*_2_, *ℳ*_3_) overestimated PP values. No difference was observed between the different definitions of lifestyle variables for PP. DP highlighted some differences among the models, including different sets of lifestyle variables. These showed an improvement in reducing bias when going from raw lifestyle variables (*ℳ*_1_, *ℳ*_11_, *ℳ*_21_) to ‘predicted’ (*ℳ*_2_, *ℳ*_12_, *ℳ*_22_) to adjusted lifestyle variables (*ℳ*_3_, *ℳ*_13_, *ℳ*_23_), decreasing the intercept estimates by up to 8 units. Similarly, SP highlighted models with ‘predicted’ lifestyle variables (*ℳ*_2_, *ℳ*_12_, *ℳ*_22_) as the best performing, although improvements were smaller. For PP, the pattern was different, with *ℳ*_0_ providing an underestimation of 8 points, while *ℳ*_1_, *ℳ*_2_, *ℳ*_3_ overestimated by 12, 11 and 12 points, respectively. However, models with both additive genetic effects and lifestyle effects, with or without their interaction, gave virtually unbiased estimates, with intercept values close to 0.

In Fig 5e,d,f, the suggestive horizontal line at 1 shows the optimal value for the estimate of the regression slope. All the models provided an underestimation of DP and SP values. However, *ℳ*_0_ gave less biased predictions, with slope equal to 0.87 and 0.97 for DP and SP, respectively. *ℳ*_1_, *ℳ*_2_, *ℳ*_3_ provided biased predictions, with slope values of 0.61, 0.65 and 0.70, respectively, for DP and 0.80, 0.82 and 0.78, respectively, for SP. Bias was slightly reduced with *ℳ*_11_, *ℳ*_12_, and *ℳ*_13_, which accounted for both genetic and lifestyle effects. and yielded values of 0.66, 0.69 and 0.73, respectively, for DP and 0.83, 0.85 and 0.83, respectively, for SP. Models with the interaction term did not show any improvement over the corresponding models without such term. For PP, *ℳ*_0_ overestimated the predictions (slope equal to 1.19), while *ℳ*_1_, *ℳ*_2_, *ℳ*_3_ close to unbiased predictions (slope equal to 0.96, 0.98 and 0.97, respectively). Considering both genetic and lifestyle effects in *ℳ*_11_, *ℳ*_12_, and *ℳ*_13_, again gave slightly more biased predictions with slope values of 1.10, 1.11 and 1.10, respectively. The same results were obtained when including the genotype-by-lifestyle interaction term *ℳ*_21_, *ℳ*_22_, *ℳ*_23_.

As expected, in the random split scenario (S1 Fig.), prediction accuracies were much higher than in the Y-O scenario, highlighting the large impact of having individuals from all the cohorts in the training set. In addition, no difference in prediction accuracy was observed among the models including raw, ‘predicted’, or adjusted lifestyle variables. The predictions obtained from all the models were unbiased in terms of the slope of the regression of true phenotypes on predicted phenotypes (S2 Fig.d,e,f).

However, while bias in terms of the intercept of the regression of true phenotypes on predicted phenotypes was generally low, some differences between the models existed (S2 Fig.a,b,c). Models with only lifestyle variables tended to be the least biased for DP and PP (S2 Fig.a,c), while models including both additive genetic effects and lifestyle effects (with or without their interaction) were virtually unbiased for SP (S2 Fig.b).

Using different sets of lifestyle variables did make a difference in terms of regression intercept. Using ‘predicted’ and adjusted lifestyle variables helped reduce bias for DP and PP (S2 Fig.a,c), with the former set being the best performing for PP, while the latter set being the best performing for DP. For SP (S2 Fig.b), using the raw lifestyle variables marginally reduced bias over the other sets of lifestyle variables.

## Discussion

### Variance partitioning and predictive ability for blood pressure traits

Obtaining accurate predictions of medically relevant phenotypes is one of the main goals of precision medicine [40]. With increasing sample size and advances in statistical methodology, PGS (*i*.*e*., predictors of phenotypes based on additive genetic effects), have started to provide appreciable prediction accuracy, for at least some complex traits [9, 10, 41]. However, PGS prediction accuracy remains low for the majority of traits. In addition, genomic heritability is substantially lower than 1 for most traits, meaning that non-genetic factors (in addition to untagged genetic variation) also play an important role in determining complex trait variation [42]. In this work, we investigated the inclusion of lifestyle variables into prediction models for blood pressure traits, namely diastolic pressure, systolic pressure, and pulse pressure. These traits have several features that make them excellent models of complex traits in general to study this problem: (1) they are modulated by different non-genetic factors (*e*.*g*., age, sex, diet, smoking); (2) they have medium values of narrow sense heritability; (3) the genomic heritability is about half the narrow sense heritability; (4) genotype-by-environment interactions explain a significant amount of blood pressure trait variance [27, 43, 44].

The LMM methodology used in this study is flexible as it allows the inclusion of genetic and non-genetic effects as well as interaction effects relatively easily [19]. On the other hand, it has been shown that the LMM assumption that all the predictors, be it genetic, environmental, or their interaction, have an effect on the trait, is not well suited for the complexity of human traits. This lack of of flexibility results in lower accuracy than current state-of-the-art PGS methods [45]. However, these methods do not easily allow the inclusion of non-genetic effects. Thus, our study should be regarded as a proof of concept, with the prediction accuracy obtained here being a lower bound on the accuracy that could be obtained with more sophisticated methods.

The results showed that lifestyle variables explained a large, medium, and small (relative to the other effects) proportion of variance for DP, SP, and PP, respectively. For each trait, the estimates of the lifestyle variance component were very similar across models (*i*.*e*., different definitions of lifestyle variables as well as other effects in the model). This shows that genetic and lifestyle components can be disentangled with the availability of relevant information and a properly balanced dataset. In these conditions, the genotype-by-lifestyle interaction component can also be estimated, although it seemed to explain a small portion of the phenotypic variance for the three traits.

However, it should be noted that for PP, the interaction variance was larger than the lifestyle variance alone. Also, it should be noted that the estimates of the genotype-by-lifestyle interaction variance were lower than in [27]. This may be explained by the different methodologies used in the two studies and, especially, by the larger sample size (280K vs 98K) used in [27]. Our decision to apply stricter filtering, especially in terms of relatedness, was aimed at avoiding confounding as much as possible.

The ‘additive’ nature of the genetic and lifestyle highlighted by the variance partition was also observed in terms of prediction accuracy. For DP and SP, using only lifestyle variables resulted in higher prediction accuracy than using only genetic variants but carried more bias, both in the intercept and slope of the regression of the observed phenotypes on the predicted phenotypes. This is in contrast with the observation that the additive genetic variance was larger than the lifestyle variance in the whole datasets (*ℳ*_0_ vs *ℳ*_1_, *ℳ*_2_ and *ℳ*_3_). This suggests that the reduced residual error in the whole dataset does not always imply increased cross-validated prediction accuracy. Yet, what seemed to be a trade-off between random error reduction on one side and bias reduction on the other side was partially solved by fitting genetic and lifestyle effects in the same model. In fact, models *ℳ*_11_, *ℳ*_12_, and *ℳ*_13_ were able to explain more phenotypic variance on the whole dataset, increase prediction accuracy and reduce bias in cross-validation. All these results go in support of including both genetic and lifestyle components in the same model for more accurate and less biased prediction of blood pressure measures at older age.

PP, measured as the difference between SP and DP, showed a different pattern across the models. The models with lifestyle effects only (*ℳ*_1_, *ℳ*_2_ and *ℳ*_3_) provided lower prediction accuracy but overall lower bias, while the model with genetic effects only (*ℳ*_0_) provided slightly higher accuracy, but overall larger bias. However, for PP, the trade-off between prediction accuracy and bias was only partially solved by accounting for both genetic and lifestyle effects in the model. In fact, for PP, models *ℳ*_11_, *ℳ*_12_, and *ℳ*_13_ gave higher predictive ability and lower bias in terms of regression intercept. Yet, models *ℳ*_1_, *ℳ*_2_ and *ℳ*_3_ were still capable of better controlling bias in terms of regression slope. This difference between PP, on one side, and DP and SP, on the other, could perhaps be explained by the different genetic correlation between younger and older individuals (Fig 3). In fact, PP showed a lower genetic correlation that could partially explain the lower predictive ability of all models for this trait.

While the interaction between the genetic and lifestyle components explained some non-negligible proportion of phenotypic variance of the three traits in the whole dataset, including this effect did not substantially improve the cross-validated predictive ability of the models. This observation agrees with previous work on several traits in UK Biobank [18] and recent theoretical work [46].

As a control experiment, we compared these results to those of a cross-validation designed using RND groups, *i*.*e*. when all cohorts were represented in the training and validation sets, while individuals were allocated to these randomly. Here, the training and validation sets are designed to be strongly linked. At the genetic level, the non-unity genetic correlation between younger and older individuals, as reported in this study, does not hamper the transfer of information between RND training and validation sets as it does for the Y-O split. At the environmental level, the distribution of lifestyle variables is homogeneous among RND groups, while it is not among the Y-O training and validation sets (see Fig 1 as well as [27]). Differences among models’ performance are small or null for the RND split, indicating that the cohort part of the model tends to downplay the contribution of the genetic and lifestyle components (as well as their interaction) (S3 Table.). Yet, cross-validation designs that include validation on known genetic groups or environmental conditions are known to pose little challenge to the prediction models. Instead, designs that stratify the validation sets based on genetic or environmental coordinates appear to be a stronger test for prediction models ([47]).

In summary, while the use of genetic and lifestyle effects appears to be a promising path to better predictions, especially in the challenging scenario of predicting phenotypes of older individuals from younger individuals, our results suggested that the patterns are somewhat different across traits. Thus, further research on additional traits is needed to understand this phenomenon more thoroughly.

### Handling of the lifestyle indicators

In this study, we sought to evaluate different ways of accounting for lifestyle in modeling blood pressure traits. We achieved this by fitting different models where the lifestyle component was defined using raw, ‘predicted’, and adjusted information of the participants’ living habits. With the baseline hypothesis that genetic variation could affect lifestyle variables, this component should be removed for proper modeling. If not, the resulting genotype-environment correlation could lead to biased estimates of the variance components and poor prediction performance, especially when modeling genotype-by-lifestyle interactions. Therefore, we decomposed the variance of each lifestyle variable and removed 1) the genetic and 2) cohort (*i*.*e*., sex and age) components. Conceptually, the remainder of the variance could be allocated into a residual component (*i*.*e*., non-accounted for by the model) and in a component that could be strictly collinear with the other lifestyle variables (*e*.*g*., body mass index as determined by the dietary preferences). Therefore, we also sought to disentangle the purely residual and the ‘lifestyle on lifestyle’ components to be used for modeling the variance of the blood pressure traits.

The **LL**^⊺^ (*i*.*e*., ‘lifestyle on lifestyle’) and **EE**^⊺^ (*i*.*e*., purely residual) kernels should contain most of the information from **MM**^⊺^ (*i*.*e*., raw lifestyle), given that the lifestyle and residual variance components were, in general, the largest components in the variance decomposition of the lifestyle variables. Therefore, by constructing these kernels, we aimed to capture different lifestyle effects while removing the genetic and cohort effects. The raw lifestyle variables were not independent of each other, and this was highlighted by the eigen decomposition of the **MM**^⊺^ kernel. This showed that 9 components were enough to explain 50% of the variance of the 27 lifestyle variables. Similarly, the eigen decomposition of the **EE**^⊺^ kernel showed that 10 components were enough to explain 50% of the variance. On the other hand, using a kernel where each of the lifestyle variables was predicted by the other 26 variables resulted in lower dimensionality, highlighting the existing covariance among the lifestyle variables themselves (*i*.*e*., the covariance between variables A, B and C increases when each of them is predicted based on the other two). For instance, decomposing the **LL**^⊺^ kernel showed that 2 components explained 50% of its variance, and 13 components explained 95% of its variance.

The different definitions of the lifestyle variables were reflected in the predictive ability of the model and partially in the proportion of variance explained. However, the trend depended on the trait. For DP, there was a slight decrease in variance explained when passing from raw lifestyle variables (**MM**^⊺^) to ’lifestyle on lifestyle’ variables (**LL**^⊺^) and to their purely residual component (**EE**^⊺^). However, the opposite trend was observed for predictive ability, where the highest accuracy and lowest bias were achieved using **EE**^⊺^. On the other hand, for SP, accuracy slightly increased and bias diminished when using kernel **LL**^⊺^. Finally, all the models performed very similarly for PP. The commonality between the two methods for adjusting the lifestyle variables is that they both did not include the genetic and cohort components of the lifestyle variables, which were properly accounted for in the models that had blood pressure measures as a dependent variable. While this treatment gave virtually no differences for PP, it gave moderate differences for SP and appreciable differences for DP. The three blood pressure traits have different architecture (Fig 2), with the lifestyle component having a large impact on DP, a lower impact on SP, and a minimal impact on PP. Thus, our study shows that traits largely determined by the lifestyle component may benefit from a pre-treatment of the lifestyle variables or, conversely, may suffer more strongly from the presence of genetic variation within the lifestyle variables. However, which treatment is more beneficial depends on the trait. Thus, researchers should carefully decide how to include environmental variables in prediction models on a trait-by-trait basis.

It appeared difficult to link the differences in prediction performance to the dimensionality of the lifestyle kernels. **LL**^⊺^ showed the lowest dimensionality, yet it was not the best performing kernel across traits. Overall, prediction performance was best for **EE**^⊺^, which showed similar dimensionality to **MM**^⊺^, the worst performing kernel. The lower dimensionality of a kernel implies a lower number of parameters to estimate, which could, in principle, reduce estimation bias in favor of environment-specific effects [46]. However, this did not seem to apply here, and rather, it appears to be the removal of non-environmental variation in the kernel that allows for best predictive performance.

## Conclusion

In this study, we investigated different ways to include lifestyle information in polygenic modeling of complex traits. We showed that pre-treatment of the lifestyle variables to remove genetic and other non-lifestyle effects can be a simple and effective way to improve predictions. However, given that the presence and magnitude of the improvement depend on the trait, more research is needed to confirm our results and extend them to more traits.

## Supporting information

**S1 Table**. Variance decomposition for the 27 lifestyle variables.

**S2 Table**. Variance decomposition with different models for the 3 blood pressure traits.

**S3 Table**. Prediction performance for different models for the 3 blood pressure traits.

**S1 Fig**. Accuracy of prediction measured as the correlation coefficient (*r*) between true phenotypes and predicted phenotype within cohorts for a) diastolic pressure, b) systolic pressure and c) pulse pressure. The colors indicate the broad class of models, *i*.*e*., including genetic effects only (light blue), lifestyle effects only (green), genetic and lifestyle effects (orange), and genetic, lifestyle and interaction effects (purple).

**S2 Fig**. Bias of predictions measured as the intercept [*β*_0_; panels a), b) and c) for diastolic pressure, systolic pressure, and pulse pressure, respectively] and slope [*β*_1_; panels d), e) and f) for diastolic pressure, systolic pressure, and pulse pressure, respectively] of the regression of the true phenotypes on the predicted phenotypes within cohorts. The colors indicate the broad class of models, *i*.*e*., including genetic effects only (light blue), lifestyle effects only (green), genetic and lifestyle effects (orange), and genetic, lifestyle and interaction effects (purple). The dotted lines indicate the expected values of *β*_0_ and *β*_1_ for unbiased predictions.

## Data and code availability

The genotype and phenotype data used in our analyses are available from UK Biobank (https://www.ukbiobank.ac.uk/). Code to reproduce the analyses is available at https://github.com/morgantelab/GxE-humans.

## Acknowledgments

This research was conducted using the UK Biobank Resource under application number 62347. We thank Trudy Mackay for helpful comments on an earlier version of this manuscript. Research reported in this publication was supported by the National Institute of General Medical Sciences of the National Institutes of Health under Award Number R35GM146868 to FM. The content is solely the responsibility of the authors and does not necessarily represent the official views of the National Institutes of Health.

## References

1. Falconer DS, Mackay TFC. Introduction to Quantitative Genetics. Pearson Education; 1996.

2. Polderman TJ, Benyamin B, De Leeuw CA, Sullivan PF, Van Bochoven A, Visscher PM, et al. Meta-analysis of the heritability of human traits based on fifty years of twin studies. Nature genetics. 2015;47(7):702–709.

3. de Los Campos G, Sorensen D, Gianola D. Genomic heritability: what is it? PLoS Genetics. 2015;11(5):e1005048.

4. Yang J, Benyamin B, McEvoy BP, Gordon S, Henders AK, Nyholt DR, et al. Common SNPs explain a large proportion of the heritability for human height. Nature genetics. 2010;42(7):565–569.

5. Ge T, Chen CY, Neale BM, Sabuncu MR, Smoller JW. Phenome-wide heritability analysis of the UK Biobank. PLoS Genetics. 2017;13(4):e1006711.

6. Consortium B, Anttila V, Bulik-Sullivan B, Finucane HK, Walters RK, Bras J, et al. Analysis of shared heritability in common disorders of the brain. Science. 2018;360(6395):eaap8757.

7. Wray NR, Yang J, Hayes BJ, Price AL, Goddard ME, Visscher PM. Pitfalls of predicting complex traits from SNPs. Nature Reviews Genetics. 2013;14(7):507–515.

8. Ma Y, Zhou X. Genetic prediction of complex traits with polygenic scores: a statistical review. Trends in Genetics. 2021;37(11):995–1011.

9. Ge T, Chen CY, Ni Y, Feng YCA, Smoller JW. Polygenic prediction via Bayesian regression and continuous shrinkage priors. Nature communications. 2019;10(1):1776.

10. Privé F, Arbel J, Vilhjálmsson BJ. LDpred2: better, faster, stronger. Bioinformatics. 2020;36(22-23):5424–5431.

11. Zhang Q, Privé F, Vilhjálmsson B, Speed D. Improved genetic prediction of complex traits from individual-level data or summary statistics. Nature communications. 2021;12(1):4192.

12. Zabad S, Gravel S, Li Y. Fast and accurate Bayesian polygenic risk modeling with variational inference. The American Journal of Human Genetics. 2023;110(5):741–761.

13. Group DPPR, et al. 10-year follow-up of diabetes incidence and weight loss in the Diabetes Prevention Program Outcomes Study. The Lancet. 2009;374(9702):1677–1686.

14. Xu W, Tan L, Wang HF, Jiang T, Tan MS, Tan L, et al. Meta-analysis of modifiable risk factors for Alzheimer’s disease. Journal of Neurology, Neurosurgery & Psychiatry. 2015;86(12):1299–1306.

15. Valenzuela PL, Carrera-Bastos P, Gálvez BG, Ruiz-Hurtado G, Ordovas JM, Ruilope LM, et al. Lifestyle interventions for the prevention and treatment of hypertension. Nature Reviews Cardiology. 2021;18(4):251–275.

16. Burkett JP, Miller GW. Using the exposome to understand environmental contributors to psychiatric disorders. Neuropsychopharmacology. 2021;46(1):263.

17. Zhou X, Van Der Werf J, Carson-Chahhoud K, Ni G, McGrath J, Hyppönen E, et al. Whole-genome approach discovers novel genetic and nongenetic variance components modulated by lifestyle for cardiovascular health. Journal of the American Heart Association. 2020;9(8):e015661.

18. Zhou X, Lee SH. An integrative analysis of genomic and exposomic data for complex traits and phenotypic prediction. Scientific reports. 2021;11(1):21495.

19. Jarquín D, Crossa J, Lacaze X, Du Cheyron P, Daucourt J, Lorgeou J, et al. A reaction norm model for genomic selection using high-dimensional genomic and environmental data. Theoretical and applied genetics. 2014;127:595–607.

20. Tiezzi F, de Los Campos G, Gaddis KP, Maltecca C. Genotype by environment (climate) interaction improves genomic prediction for production traits in US Holstein cattle. Journal of dairy science. 2017;100(3):2042–2056.

21. Huang W, Carbone MA, Lyman RF, Anholt RR, Mackay TF. Genotype by environment interaction for gene expression in Drosophila melanogaster. Nature communications. 2020;11(1):5451.

22. Rogers AR, Dunne JC, Romay C, Bohn M, Buckler ES, Ciampitti IA, et al. The importance of dominance and genotype-by-environment interactions on grain yield variation in a large-scale public cooperative maize experiment. G3. 2021;11(2):jkaa050.

23. Chen SY, Freitas PH, Oliveira HR, Lázaro SF, Huang YJ, Howard JT, et al. Genotype-by-environment interactions for reproduction, body composition, and growth traits in maternal-line pigs based on single-step genomic reaction norms. Genetics Selection Evolution. 2021;53(1):51.

24. Napier JD, Heckman RW, Juenger TE. Gene-by-environment interactions in plants: Molecular mechanisms, environmental drivers, and adaptive plasticity. The Plant Cell. 2023;35(1):109–124.

25. de Leeuw CA, Stringer S, Dekkers IA, Heskes T, Posthuma D. Conditional and interaction gene-set analysis reveals novel functional pathways for blood pressure. Nature communications. 2018;9(1):3768.

26. Moore R, Casale FP, Jan Bonder M, Horta D, Franke L, Barroso I, et al. A linear mixed-model approach to study multivariate gene–environment interactions. Nature genetics. 2019;51(1):180–186.

27. Kerin M, Marchini J. Inferring gene-by-environment interactions with a Bayesian whole-genome regression model. The American Journal of Human Genetics. 2020;107(4):698–713.

28. Laville V, Majarian T, Sung YJ, Schwander K, Feitosa MF, Chasman DI, et al. Gene-lifestyle interactions in the genomics of human complex traits. European Journal of Human Genetics. 2022;30(6):730–739.

29. Jung HU, Kim DJ, Baek EJ, Chung JY, Ha TW, Kim HK, et al. Gene-environment interaction explains a part of missing heritability in human body mass index. Communications Biology. 2023;6(1):324.

30. Durvasula A, Price A. Distinct explanations underlie gene-environment interactions in the UK Biobank. medRxiv. 2023; p. 2023–09.

31. Jaffee SR, Price TS. Gene–environment correlations: A review of the evidence and implications for prevention of mental illness. Molecular psychiatry. 2007;12(5):432–442.

32. Bycroft C, Freeman C, Petkova D, Band G, Elliott LT, Sharp K, et al. The UK Biobank resource with deep phenotyping and genomic data. Nature. 2018;562(7726):203–209.

33. R Core Team. R: A Language and Environment for Statistical Computing; 2023. Available from: https://www.R-project.org/.

34. Wickham H. ggplot2: Elegant Graphics for Data Analysis. Springer-Verlag New York; 2016. Available from: https://ggplot2.tidyverse.org.

35. Tobin MD, Sheehan NA, Scurrah KJ, Burton PR. Adjusting for treatment effects in studies of quantitative traits: antihypertensive therapy and systolic blood pressure. Statistics in medicine. 2005;24(19):2911–2935.

36. Yang J, Lee SH, Goddard ME, Visscher PM. GCTA: a tool for genome-wide complex trait analysis. The American Journal of Human Genetics. 2011;88(1):76–82.

37. Pérez P, de Los Campos G. Genome-wide regression and prediction with the BGLR statistical package. Genetics. 2014;198(2):483–495.

38. Snow G. TeachingDemos: Demonstrations for Teaching and Learning; 2024. Available from: https://CRAN.R-project.org/package=TeachingDemos.

39. Legarra A, Reverter A. Semi-parametric estimates of population accuracy and bias of predictions of breeding values and future phenotypes using the LR method. Genetics Selection Evolution. 2018;50:1–18.

40. Lewis CM, Vassos E. Polygenic risk scores: from research tools to clinical instruments. Genome medicine. 2020;12(1):44.

41. Khera AV, Chaffin M, Aragam KG, Haas ME, Roselli C, Choi SH, et al. Genome-wide polygenic scores for common diseases identify individuals with risk equivalent to monogenic mutations. Nature genetics. 2018;50(9):1219–1224.

42. Canela-Xandri O, Rawlik K, Tenesa A. An atlas of genetic associations in UK Biobank. Nature genetics. 2018;50(11):1593–1599.

43. Waken R, de Las Fuentes L, Rao D. A review of the genetics of hypertension with a focus on gene-environment interactions. Current hypertension reports. 2017;19:1–8.

44. Evangelou E, Warren HR, Mosen-Ansorena D, Mifsud B, Pazoki R, Gao H, et al. Genetic analysis of over 1 million people identifies 535 new loci associated with blood pressure traits. Nature genetics. 2018;50(10):1412–1425.

45. Lloyd-Jones LR, Zeng J, Sidorenko J, Yengo L, Moser G, Kemper KE, et al. Improved polygenic prediction by Bayesian multiple regression on summary statistics. Nature communications. 2019;10(1):5086.

46. Weine E, Smith SP, Knowlton RK, Harpak A. Tradeoffs in Modeling Context Dependency in Complex Trait Genetics. bioRxiv. 2023; p. 2023–06.

47. Tiezzi F, Fleming A, Malchiodi F. Use of Milk Infrared Spectral Data as Environmental Covariates in Genomic Prediction Models for Production Traits in Canadian Holstein. Animals. 2022;12(9):1189.

