## Supplementary figures and images for "Using lifestyle information in polygenic modeling of blood pressure traits: a simple method to reduce bias"

### S1 Fig

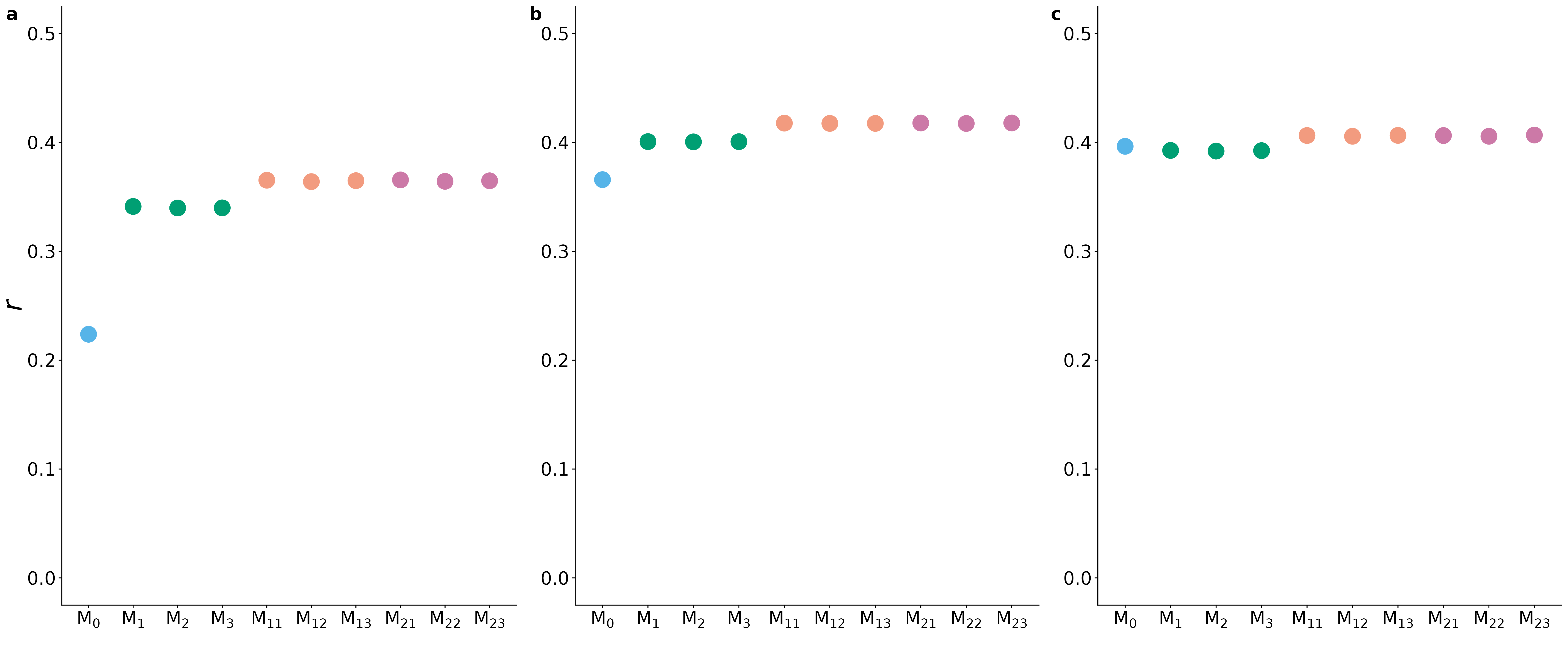

### S2 Fig

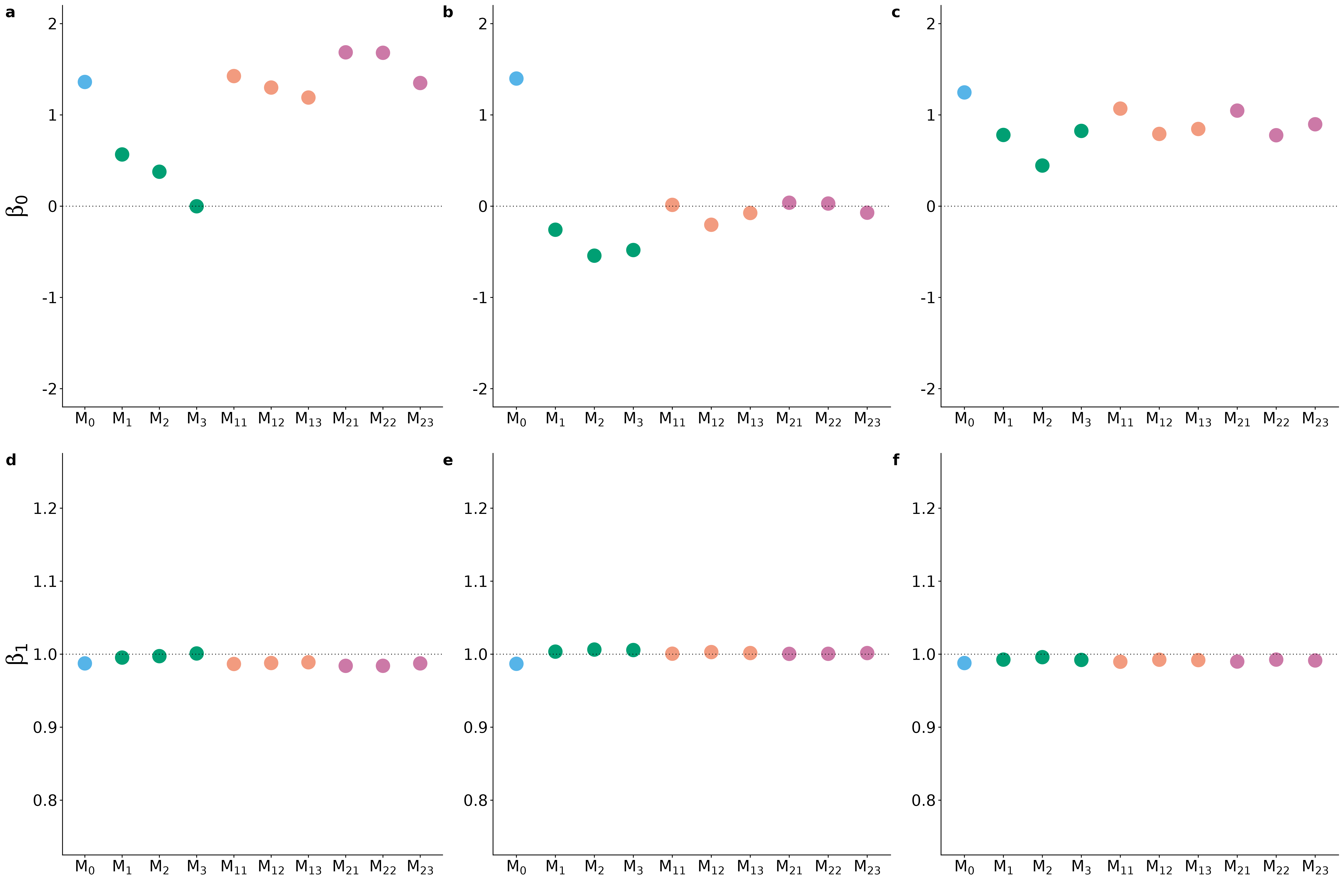
